# Hidden Markov Modeling of Molecular Orientations and Structures from High-Speed Atomic Force Microscopy Time-Series Images

**DOI:** 10.1101/2022.07.18.500393

**Authors:** Tomonori Ogane, Daisuke Noshiro, Toshio Ando, Atsuko Yamashita, Yuji Sugita, Yasuhiro Matsunaga

## Abstract

High-speed atomic force microscopy (HS-AFM) is a powerful technique for capturing the time-resolved behavior of biomolecules. However, structural information in HS-AFM images is limited to the surface geometry of a sample molecule. Inferring latent three-dimensional structures from the surface geometry is thus important for getting more insights into conformational dynamics of a target biomolecule. Existing methods for estimating the structures are based on the rigid-body fitting of candidate structures to each frame of HS-AFM images. Here, we extend the existing frame-by-frame rigid-body fitting analysis to multiple frames to exploit orientational correlations of a sample molecule between adjacent frames in HS-AFM data due to the interaction with the stage. In the method, we treat HS-AFM data as time-series data, and they are analyzed with the hidden Markov modeling. Using simulated HS-AFM images of the taste receptor type 1 as a test case, the proposed method shows a more robust estimation of molecular orientations than the frame-by-frame analysis. The method is widely applicable in integrative modeling of conformational dynamics using real experimental HS-AFM data.

**Author summary:** Biomolecules dynamically change their structures in cells and perform various functions that are important for life. It is difficult to observe such dynamic structural changes using conventional experimental techniques, such as X-ray crystallography, that mainly observe static atomic structures. Alternative approach is the atomic force microscopy (AFM), a technique to measure the surface geometry of a sample molecule by scanning with an acute tip. Recently, by increasing the imaging rate of the AFM, it has become possible to observe the dynamics of biomolecules at work. However, the information measured with AFM is limited to the surface geometry of a molecule, so it is important to extract three-dimensional structural information from those data. In this study, we develop a method to estimate structural information with higher accuracy.

## Introduction

Observing conformational dynamics of biomolecules in action is important to deepen our understanding of biomolecular functions. Among the various types of experimental measurements, single-molecule measurements are powerful approaches since they can directly characterize heterogeneous fluctuations of biomolecules. The measurement techniques used in single-molecule measurements include Förster resonance energy transfer microscopies (FRET) [1][2], optical tweezers [3], and atomic force microscopy (AFM) [4]. AFM uses an acute tip as a probe for scanning the surface of a molecule and the information on the surface morphology can be obtained. In general, the imaging rate of conventional AFM is too slow to observe conformational dynamics of biomolecule in action. To overcome this limitation, the high-speed AFM (HS-AFM) have been developed by improving the resonant frequency of cantilevers, the speed of scanners [5][6]. HS-AFM enables us to observe the direct visualization of biomolecules in action at high spatiotemporal resolution. Indeed, HS-AFM has been widely applied to visualize conformational dynamics of various biomolecules; for example, myosin V walking [7], rotary catalysis of F1-ATPase [8], the conformational dynamics of CRISPR-Cas9 in action [9], and the structure and dynamics of intrinsically disordered proteins [10].

A weak point of AFM is that the observed information is limited to the geometry on the molecular surface. Also, the obtained image profile depends on the tip shape as well as the molecular surface. Therefore, estimating the real 3D structural information hidden in the 2D AFM data is an important remaining problem. In HS-AFM and AFM observations, molecules tend to be fixed in specific orientations due to interactions with the stage, and the observed data are anisotropic. Generally, it is impossible to reconstruct structure from anisotropic data in a model-free way as is done in the single particle analysis of cryo-electron microscopy (CryoEM) in which a tremendous number of isotropic images are taken. Thus, a model-based approach is required to infer 3D structures from HS-AFM and AFM data. In this direction, the most popular model-based approach is the rigid-body fitting method [11][12][13][14][15][16][17][18][19][20]. In this method, assuming that target molecules in AFM have known structures already observed by X-ray, NMR, or CryoEM, many candidate structures are superimposed to AFM images to determine the best structure matching with the AFM image. The number of candidate structures is increased by, for example, performing molecular dynamics (MD) simulations using an experimental structure as an initial condition.

Niina et al. recently treated a candidate structural model as a rigid-body, and by exhaustively rotating and translating it to generate a pseudo AFM image and superimposing it on the HS-AFM image [19]. They successfully estimated the appropriate position and orientation of the molecule. Moreover, by repeating this calculation using various tip radii and half angles (see Fig. 1A), these parameters were also successfully estimated. In another study [21], combined with MD simulations, a flexible fitting method was developed to search for the optimal structure by changing the structure to fit the AFM image, which enables estimation of the 3D structure even when there is no “correct” candidate structure. However, a drawback of the flexible fitting is its high computational cost, which makes it difficult to apply to a series of images obtained by HS-AFM.

**Figure 1.**
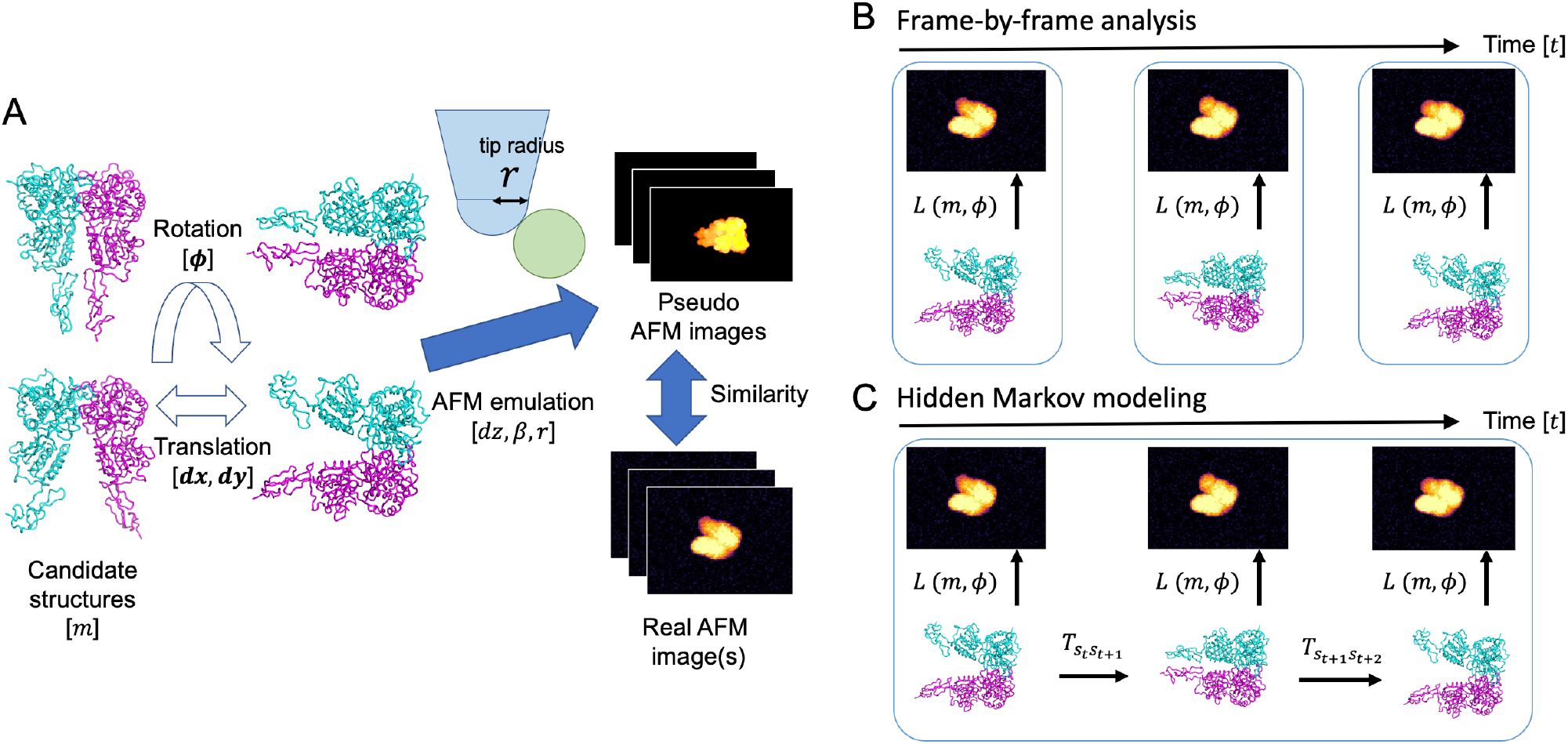
Scheme of analysis on atomic force microscopy images. (A) Computation of likelihood of each atomic force microcopy image. A structure is rotated and translated, and then emulated to generate a pseudo image. The pseudo image is compared with real images with some similarity measures. (B) Schemes of the frame-by-frame rigid body fitting. (C) Schematic of hidden Markov modeling using multiple images as time-series data.

Usually, the rigid-body fitting and flexible fitting are based on the idea that the 3D structure is independently estimated by frame-by-frame analysis of each image. On the other hand, a unique point of HS-AFM data is that they are time-series data where adjacent frames are temporally correlated with other. As mentioned above, because biomolecules interact with the stage, the orientations of the molecules tend to be fixed in a particular direction, thus orientational correlations exist even in the millisecond to second time-scale. Focusing on this unique feature of HS-AFM time series data, Fuchigami et al recently applied a time-series analysis method, called the particle filter [22] to artificially simulated AFM images. In the particle filter, a large number of copies (or replicas) is simultaneously simulated in MD, and the copies are duplicated or deleted according to the likelihood of the corresponding image. By repeating this process through the whole image frames, one can obtain a sequence of structures that well matches with time-series of AFM images. The particle filter is based on the state-space modeling. It assumes that experimental data are generated by projection from a hidden state space, using the three-dimensional structure space of a target molecule as a state space. It is theoretically guaranteed that a Bayesian inference (posterior density) of the latent space is obtained by the computational algorithm of the particle filter. The drawback of the particle filter, however, is that it is computationally very expensive. In fact, a number of simulation copies (replicas) required to accurately sample the posterior densities generally increases exponentially with the state space dimension.

This study develops a method to treat multiple HS-AFM images together as time-series data for more accurate structure and orientation estimations, while keeping computational cost lower than the particle filter. Similar to the particle filter, our idea is based on the use of the 3D structure and orientation space as a state space, but instead of a continuous space, we use a discretized space by clustering the candidate structures. By using a discretized state space model, we can apply an efficient estimation algorithm based on hidden Markov modeling [23] instead of the computationally expensive MD simulations. We combine our method with the existing rigid-body fitting method and show that our hidden Markov modeling is a natural extension of the frame-by-frame rigid-body fitting. By performing twin experiments, we see that our method can more accurately estimate the orientation of the structure than the existing rigid-body fitting method. Furthermore, we show that our estimation method is robust even when the geometry of the tip shape in modeling differs from the correct (ground-truth) one in experiment. Finally, we discuss the applicability of the method for integrative modeling on conformational dynamics to real experimental HS-AFM data.

## Methods

### Frame-by-frame rigid-body fitting

This study proposes a method to analyze multiple AFM images as a time series by extending the conventional rigid-body fitting method. First, we review the conventional rigid-body fitting, in which each AFM image is analyzed by a frame-by-frame manner, and then we describe our hidden Markov approach in the next subsection.

In the rigid-body fitting, an ensemble of candidate structures is prepared from experimentally determined structures or structures obtained by MD simulations. The goal of rigid-body fitting is to select the correct structure and to find its orientation and translation from a given experimental AFM image. To accomplish this, the rigid-body fitting generates a large number of synthetic AFM images with various structures *m*, 3D rotations *ϕ* (quaternions are used in this study), translations (*dx, dy*), and other parameters, and compares them to the experimental AFM images. The parameter set that produces the synthetic AFM image with the highest similarity to the experimental AFM image is considered as the best estimate. Hereafter, we will refer to these synthetic images as pseudo-AFM images.

To generate pseudo-AFM images, we employed a conventional collision-detection method [11][18][19]. This method calculates the height at which a tip perpendicular to the stage collides with a sample molecule. Let the stage be the *xy*-plane, then the height of the stage (*z*-coordinate) is aligned with the minimum value of the *z*-coordinate of the atoms of the molecule. Then, let us describe by *dz*an offset of the stage from the origin as an unknown parameter. The scaling parameter *β*for height *z* is also considered as unknown, if the calibration of the height scale is not perfect. Following Niina et al. [19], we approximate the tip by a hemisphere (with radius *r*) combined with a circular frustum of a cone. In summary, unknown parameters when generating pseudo-AFM images are structure *m*, rotation *ϕ*, and translation (*dx, dy*), offset *dz*, scaling parameter *β*to calibrate the height, and apex radius *r* of the tip.

Rigid-body fitting measures similarity between the generated pseudo-AFM image and the experimental AFM image. In this approach, the parameters used to generate the pseudo-AFM image with the highest similarity are considered as the best estimate. The similarity measures include pixel-RMSD [22], cross correlation [18][20], cosine similarity [19], and structural similarity index measure (SSIM) [17]. For example, pixel-RMSD is defined as follows, essentially the same as peak signal-to-noise ratio (PSNR):

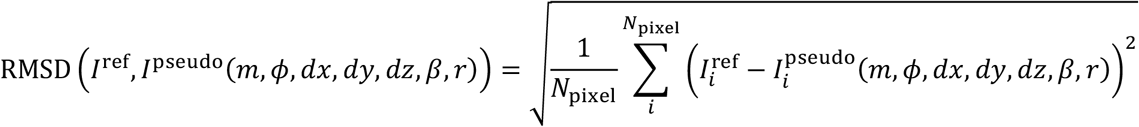

where 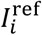 and 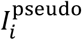 are the heights at pixel *i* of the experimental AFM and the pseudo-AFM images, respectively. *N*_pixel_ is the number of pixels in the image. In this case, a smaller pixel-RMSD means a higher similarity.

Here, we introduce the concept of probability into the rigid-body fitting to extend the analysis to a likelihood-based analysis that allows more flexible modeling. First, we assume that an experimental AFM image can be generated by adding spatially independent Gaussian noise to a pseudo-AFM image. Then, the probability of observing the experimental AFM image or likelihood, given a set of parameters, can be written as follows

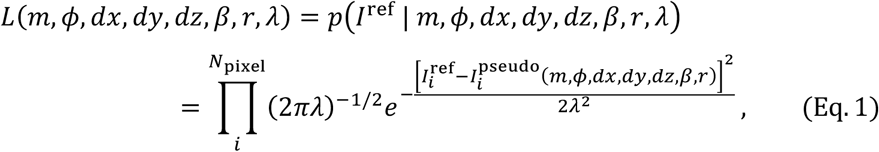

where *λ* is the standard deviation of the Gaussian noise. From the above equation, it is obvious that this likelihood *L* is maximized for the parameter set with the smallest pixel-RMSD. In principle, the maximum likelihood parameters can be found by exhaustively searching possible parameters, but the large number of possible combinations of the parameters makes it infeasible to search all of them. Therefore, in this study, we reduce the number of parameters by integrating out some of unknown parameters, as was done by Cossio and Hummer in their analysis of CyoEM images [24]. In Eq. 1, the parameters, *dz, β*, and *λ*, can be analytically eliminated by integration them out. Then, the marginalized likelihood, as derived by Cossio and Hummer, becomes as follows,

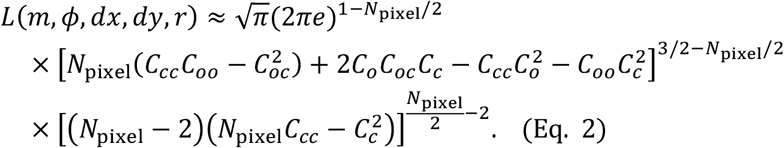

Here, *N*_pixel_ is the total number of pixels in the image. *C*_*o*_, *C*_*c*_, *C*_*oo*_, *C*_*cc*_, *C*_*oc*_ are sums of pixel heights, sums of squares, and inner product of pixel heights, respectively:

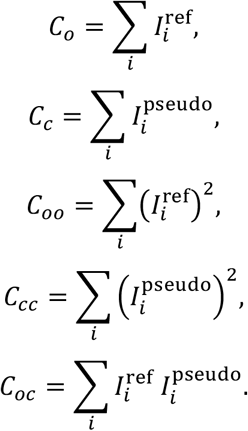

This marginalized likelihood depends only on the candidate structure *m*, 3D rotation *ϕ*, translation (*dx, dy*) and tip apex radius *r*, and now this is more feasible compared to Eq. 1.

In this study, using the marginalized likelihood (Eq. 2), we conducted maximum likelihood estimates for the candidate structure *m*, 3D rotation *ϕ*, and tip apex radius *r*. For the integration over the 3D rotation *ϕ*, we used 577 quaternions which are uniformly sampled in the SO(3) group space [25][26]. *C*_*o*_, *C*_*c*_, *C*_*oo*_ and *C*_*cc*_ are invariant with respect to translation (*dx, dy*), thus only *C*_*oc*_ is needed to integrate over (*dx, dy*). *C*_*oc*_ can be expressed in the form of convolution integrals, thus can be efficiently computed in the inverse space using Fast Fourier Transform (FFT) [26]. In this study, we employed FFT and the computational cost was reduced from 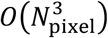 to 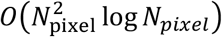.

### Hidden Markov modeling extension to multiple frames

To extend the rigid-body fitting described in the previous subsection to treat multiple AFM images as a time-series, we introduce a likelihood for multiple images consisting of temporally consecutive frames (Fig. 1C). To define this, we first define state *s* (*s* = 1, …, *M*) by the combination of the candidate structure *m* and the 3D rotation *ϕ*, and then introduce the transition probability from state *s*_G_ at the *t*-th frame (*t* = 1, …, *T*) to state *s*_GHO_ at the (*t* + 1)-th frame as 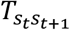. We assume that the transitions between states satisfy the Markovian property so that the transitions can be fully characterized by only the transition probabilities. Furthermore, the transition probabilities are assumed to satisfy the detailed balance. Translations (*dx, dy*) are not included in the definition of the state because, unlike rotation, translations may not affect the conformational dynamics of a molecule because interactions with the stage do not change over translations. Then, the probability of being in state *s*_1_ at the 1st frame and observing image *I*^*ref*,1^ can be written by *p*(*s*_1_)*L*(*m*_1_, *ϕ* _1_, *dx*_1_, *dy*_1_, *r*) where *m*_1_, *ϕ* _1_, *dx*_1_, *dy*_1_ are the parameter sets at the 1st frame, *p*(*s*_1_) is the equilibrium probability of *s*_1_. Repeating this, the probability of observing state *s*_2_ and image *I*^*ref*,2^ in the 2nd frame is written by 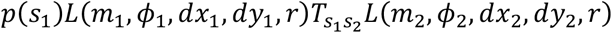 using the transition probability. By repeating this process up to the *T*th frame, and marginalizing over the state *s*_G_ for all the frames, the likelihood of AFM time-series images becomes as follows,

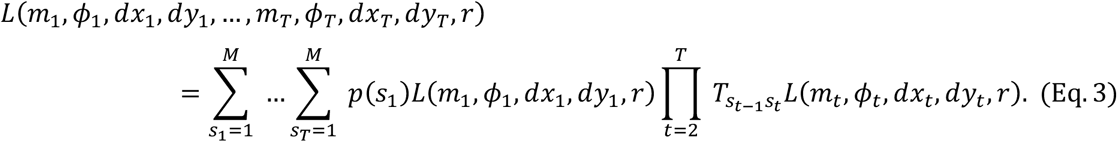

Note that if the equilibrium probabilities and transition probabilities are uniform, the likelihood in Eq. 3 is equivalent to simply multiplying the likelihoods of frame-by-frame rigid-body fitting. On the other hand, if the transition probabilities are not uniform, the likelihood is affected by not only the individual likelihood of rigid-body fitting, but also the transition probabilities.

Since the transition probabilities 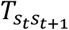 are usually unknown in advance of the analysis of AFM images, they should be estimated from the data using the Baum-Welch algorithm of hidden Markov modeling [23]. In the original Baum-Welch algorithm, the so-called *emission*, which is the probability of observing an image from a given state, is estimated together from the data. Since the emission is exactly the same as the likelihood *L*(*m, ϕ, d*x, *dy, r*) of the frame-by-frame rigid-body fitting in this case, only the transition probabilities are estimated here. The Baum-Welch algorithm can estimate transition probabilities that reproduce the data with higher likelihood defined in Eq. 3 (though it only locally optimizes the likelihood thus it could be trapped a local minimum). In the algorithm, the Forward-Backward algorithm is used for the calculation of the likelihood, which drastically reduces the computational cost from *O*(2*TM*^*T*^) to *O*(2*TM*^2^).

An important characteristic of the Baum-Welch algorithm is that the transition probability, initially sets to zero, remains zero even after the algorithm is applied [23]. By imposing these topological constraints as initial conditions, it is possible to introduce a prior knowledge that transitions between certain states completely unlikely happen in a single transition and further reduce computational cost [27]. In the case of AFM experiments, a target molecule is often interacting with the stage, and the probability of large rotations between adjacent frames does not likely happen. Thus, we propose to impose a prior knowledge that significantly change its orientations unlikely occur in a single transition. This can be achieved by imposing the transition probabilities to be zero between states with largely different orientations. Throughout this study, we imposed the probability of transitions between states whose quaternion distance is 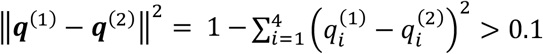 to be zero.

In the calculation of the likelihood of AFM time-series images (Eq. 3), we first need to compute the likelihood *L*(*m*_G_, *ϕ* _G_, *dx*_G_, *dy*_G_, *r*) of the frame-by-frame rigid-body fitting (Eq. 2). The results of the frame-by-frame likelihood can be used to further reduce the number of states used in Eq. 3, because the structures and orientations with very small frame-by-frame likelihood (Eq. 2) value are also very small even in the likelihood of the time-series (Eq. 3). By removing the structures and orientations with very small frame-by-frame likelihoods, we can further reduce computational cost of the hidden Markov modeling.

Once the transition probabilities are estimated by the Baum-Welch algorithm, they can be used to estimate the maximum likelihood states that match the AFM time-series images. This maximum likelihood estimation can be done efficiently with the Viterbi algorithm in the Hidden Markov modeling [23].

### Twin Experiment

To compare the performance of the Hidden Markov modeling method described in the previous subsection with the frame-by-frame rigid-body fitting, we performed twin experiments. In the experiments, the accuracy of the method is assessed by generating artificial “experimental” AFM images and compare the estimated structures and orientations with the ground-truths used to generate the images.

We used the extracellular ligand-binding domains (LBDs) of the medaka fish taste receptor (PDB ID: 5X2M [28]) as a molecular model in the twin experiments (Fig. 2A). This is a heterodimer consisting of T1r2a and T1r3. The ligand-binding domains (LBDs) in T1r2a and T1r3 have a ligand-binding pocket. Previous biophysical studies have shown that the T1r2aLBD pocket is characterized by a broad yet discriminating chemical recognition, while the T1r3LBD pocket is rather loosely bound an amino acid [28]. The receptor is an ortholog of human sweet and umami taste receptors, in which each subunit of the heterodimer is known to play different functional roles [29]. In this context, it is important to correctly identify T1r2aLBD and T1r3LBD of this pseudo symmetric structure from AFM images and to characterize the structure and dynamics of each domain.

**Figure 2.**
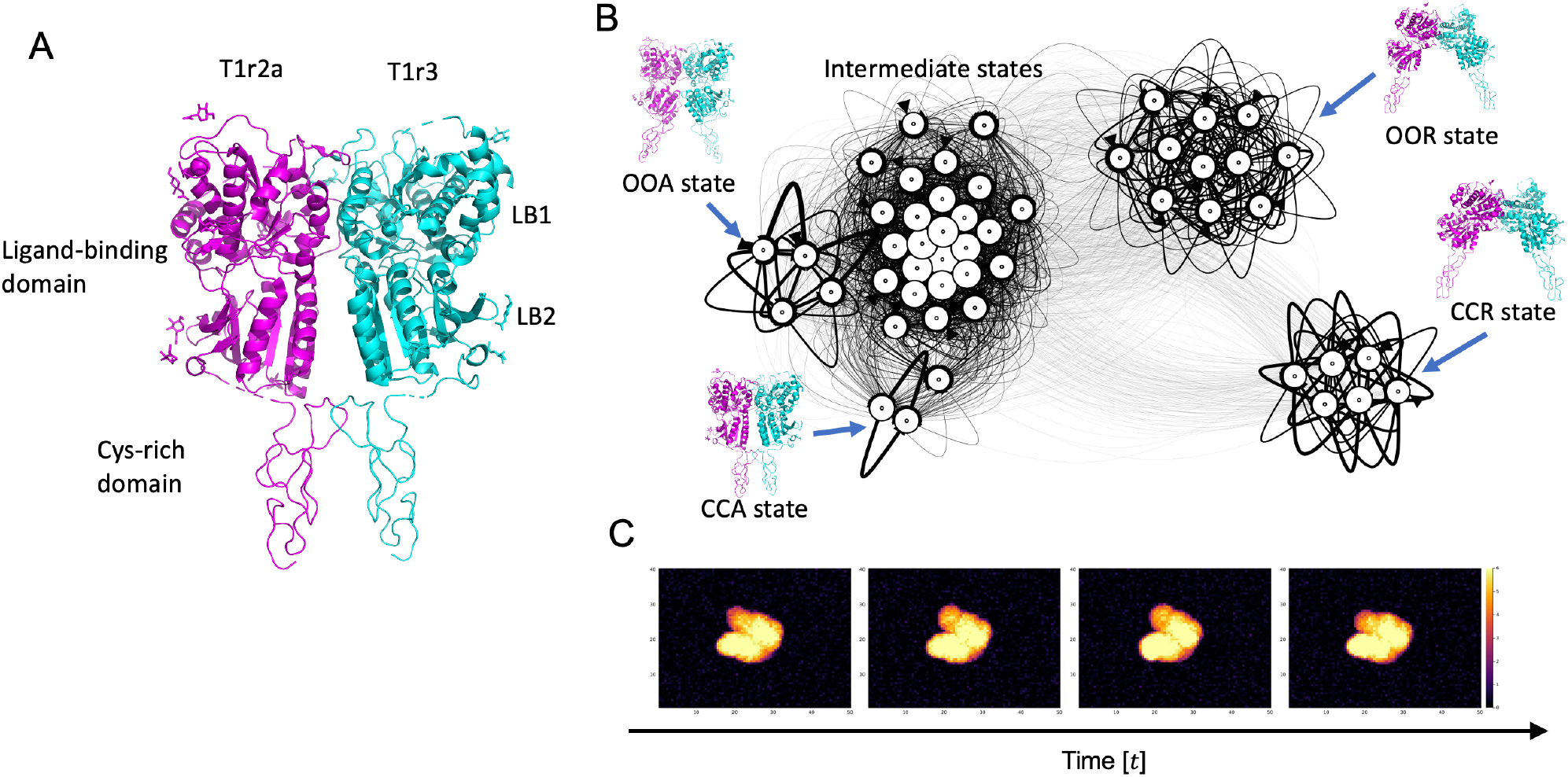
Structure of the taste receptor and its Markov state model (MSM). (A) The extracellular ligand-binding domains and the cysteine-rich domains of the medaka fish taste receptor. (B) MSM constructed from coarse-grained simulation data. (C) “Experimental” AFM data generated by a stochastic simulation with the MSM, which is used for the twin experiments.

To generate artificial “experimental” AFM images in the twin experiments, we performed coarse-grained MD simulations of the taste receptor (without the stage of AFM), sampling all possible structures, and then constructed a Markov state model (MSM) [30][31][32]. Before starting MD simulations, to clarify the dynamics of the entire extracellular region not only with LBD but also with the downstream cysteine-rich domain, cysteine-rich domains were added to the atomistic structure of 5X2M by homology modeling using the structure of the same protein family with T1rs, class C G protein-coupled receptor (GPCR), the active form of human calcium-sensing receptor extracellular domain (PDB ID: 5K5S [33]) as a template. Then, the subdomains (LB1 and LB2) of LBDs were superimposed on the structure of the inactive form of human calcium-sensing receptor extracellular domain (PDB ID: 5K5T [33]) to create a structure where LBDs are open (called the OO state while the closed state is called the CC state). Also, cysteine-rich domain was also superimposed on the structure of a single protomer of another class C GPCR, the metabotropic receptor (mGluR)3LBD (PDB ID: 2E4U [34]), to create an open structure of the whole dimer (called the R state while the closed state is called the A state). Simple superposition of the LBDs made the interface region of the LBDs sterically crashed, so the structure was superimposed again to the crashed structure by a targeted MD simulation [35] using an all-atom model from the 5X2M structure. In the targeted MD simulation, we did not impose any pulling forces to the interface region to make the regions relaxed according to naturally occurring inter-molecular interactions. Similar calculations were performed for structures with open LBDs (OO state). All obtained structures were then subjected to all-atom MD with positional restraints on their C*α* atoms to relax the other regions including side-chains and surrounding solvents. Then, using the obtained structures, C*α*-based Karanicolas-Brooks (KS) Go-model [36][37] was created for coarse-grained MD simulations. Based on KB Go-model, we created a quad-basin potential energy by macro-mixing the four states (CCA, CCR, OOR, OOA states, see Fig. 2B) obtained by the all-atom modeling, which makes its energy landscape stabilizing these four states. Then, a number of coarse-grained MD simulations were conducted with the created model to sample all possible structures.

All the MD simulations were performed with GENESIS [38][39]. In the all-atom MD simulations, the simulation systems were prepared with the CHARMM-GUI web server [40] using the modeled structures. Missing residues were modelled with MODELLER [41]. CHARMM36 force field [42][43] was used with the TIP3P water model [44]. Electrostatic interactions were treated using the Smooth Particle Mesh Ewald method [45], and covalent bonds including hydrogen atoms were constrained using SHAKE [46] and SETTLE [47]. Temperature was kept at 303.15 K with Langevin dynamics. In the coarse-grained MD simulations, the parameter sets for KB Go-model were prepared by using the MMTSB [48] Go model web server. Quad-basin macro-mixing parameters were created from four single-basin KB Go-model parameters. All the bonds between coarse-grained atoms were constrained using SHAKE, and the temperature was controlled at 200 K by Langevin dynamics.

The obtained MD structures were aligned to the CCA structure, and principal component analysis (PCA [49]) was performed on the Cartesian coordinates of the aligned structures. The 1st PC corresponds to opening-closing structural motions in the entire dimer, and the 2nd PC corresponds to opening-closing motions in LB1 and LB2 of LBDs. We performed the *k*-means clustering in the space of these 1st and 2nd PCs to define the states in the Markov state model (MSM) with *k* = 50. Note that, “states” used here, contain only structures (does not contain rotations). By counting the transition between these states from the trajectories, transition probabilities for states (structures) were estimated. Finally, we constructed a MSM of the taste receptor to represent its conformational dynamics.

We used the constructed MSM to generate artificial “experimental” AFM data. To generate AFM images (Fig. S1), (i) stochastic simulations were performed with the MSM to calculate the transitions between states for 100 frames. According to the obtained sequence of states, the sequence of corresponding structures was drawn from the centroid of *k*-means clustering. Then, (ii) 100 frames of AFM images were created from the sequence of structures using a collision-detection method. (iii) To mimic experimental noises, spatially independent Gaussian noise with a standard deviation of 0.3 nm were added to the generated AFM images. The standard deviation of 0.3 nm is a typical noise width of AFM images [19]. The pixel resolution is 0.625 nm × 0.625 nm, and the image has 80 pixels along the X-axis and 60 pixels along the Y-axis. When calculating the collision with a tip, the effective radius of the C*α* atom for its amino acid residue was taken from the CryoEM study [24] while a radius of 2.5 nm was used for a tip. For simplicity, we assumed that a molecule is strongly interacted with the stage, so the orientation of the molecule is fixed to a specific angle during the 100 frames. To determine the orientation, a quaternion was randomly selected from 576 quaternions uniformly sampled in SO(3) group [25][26]. Irrespective the chosen orientations, the same transition probabilities between states (structures) were used for the stochastic simulation. The above calculations were repeated 50 times to produce a total of 50 sets of 100-frame AFM data.

We performed the frame-by-frame rigid-body fitting and the maximum likelihood estimation with hidden Markov modeling on the “experimental” AFM images created by the above procedure. We used the same 50 structures with the MSM (i.e., the centroids of *k*-means clustering) as candidate structures. The same 576 quaternions and AFM image grids were used. RMSD between the ground-truth and estimated structures was used to access the accuracy of the estimation. In order to access the accuracy of not only the structure *m* but also the 3D rotation *ϕ*, the RMSD calculation was performed without applying structural alignment.

The actual experimental data often suggest that the shape of a tip is not simple as a circular frustum of a cone assumed in this study, but a more complex one. Therefore, to account for the possibility that the shape of the real tip might be different from the assumed one, we performed maximum likelihood estimations with intentionally different tip apex radii (ground truth used for generating “experimental” images are 2.5 nm while we used 1.5 nm, 1.8 nm, 2.5 nm, 3.2 nm, and 3.5 nm for estimation). We tested how robustly the method estimates the structure and orientation in the situation where there was a discrepancy in the tip apex radius. Furthermore, the hidden Markov modeling can estimate the transition probabilities 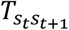 between states. We reconstructed the transition probabilities between structures by reducing orientations from 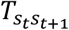obtained by the Baum-Welch algorithm. Then, compared the reduced transition probabilities between structures with those of the Markov state model.

## Software availability

The Hidden Markov modeling-based analysis method described here is implemented and publicly available as part of the MDToolbox.jl package and can be downloaded from https://github.com/matsunagalab/MDToolbox.jl. In addition, Jupyter notebooks to reproduce the results of this paper are publicly available at https://github.com/matsunagalab/paper_ogane2022.

## Results and Discussion

### Twin experiments with identical radii

First, we examined the estimation accuracy when the ground-truth tip radius used to generate the experimental AFM data (2.5 nm) and the tip radius used for the maximum likelihood estimation are identical. Figure 3C shows that both methods (the frame-by-frame rigid-body fitting and the hidden Markov modeling) are able to estimate the correct ones from the combinations of 50 candidate structures and 576 angles with almost 100 % accuracy, despite the addition of Gaussian noise with a standard deviation of 0.3 nm. Specifically, the frame-by-frame rigid body fitting resulted in zero RMSD (without structural alignment) for 4991 frames out of 50 frames × 100 sets of images. The Hidden Markov modeling resulted in zero RMSD in the same number of frames (4991 frames). The accuracy of this estimation result may be dependent on the noise size; in the current setup, the spatial scale of the average RMSD (∼1.1 nm) between the 50 structures and the resolution of the rotations is comparable or smaller than the size of Gaussian noise.

**Figure 3.**
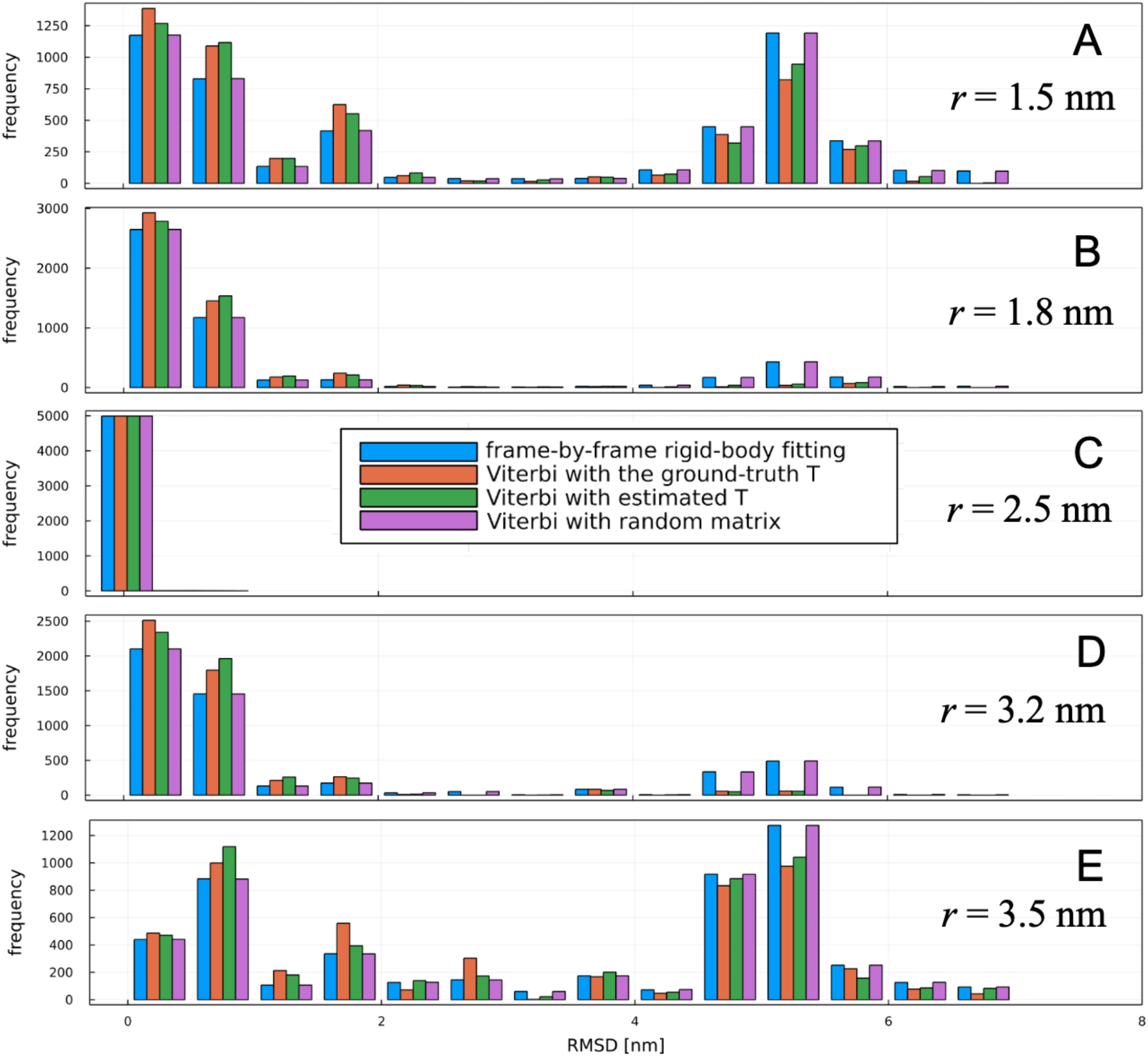
Results of twin experiments. Histograms of root-mean square deviations (RMSDs) of estimated structures from the ground-truth structures in twin experiments. Note that the RMSDs were computed without structural alignment. Structural estimations were performed with various conditions: with different algorithms (the frame-by-frame rigid-body fitting, the Viterbi algorithm using the ground-truth transition probabilities, estimated probabilities by the Baum-Welch algorithm, and a random matrix), and different tip radii (1.5 nm, 1.8 nm, 2.5 nm that is the ground-truth, 3.2 nm, 3.5 nm).

### Twin experiments with discrepant radii

Next, we examined the estimation accuracy when the ground-truth tip radius (2.5 nm) used in generating the AFM data is different from the tip radius used for the maximum likelihood estimation. As described in Methods, the actual experimental probe geometry is more complex than the assumed shape, so we intentionally used a different radius for the estimation to assess how the estimations are robust against such discrepancies.

Figures 3A, 3B, 3D, and 3E show the accuracies of the frame-by-frame rigid-body fitting with different tip radii, measured by structural RMSD from the ground-truth structure. When the radius difference is close, the RMSDs are distributed lower than 1.0 nm. However, as the radius difference increases, higher peaks appear at around the RMSD of 5.0 nm, indicating that the estimation accuracy is becoming worse. The structures around the RMSD of 5.0 nm are flipped 180 degrees from the ground-truth structure, indicating that the estimations failed to assign each monomer of the taste receptor to the AFM image correctly. This is because, as the radius difference increases, the likelihood fails to catch the subtle surface differences in the two monomers.

To further investigate the cause of the double peaks in the RMSD distribution, we examined the the relation between the orientations and the likelihood values (Eq. 2) using the first frame of the first set in the AFM data (shown in Fig. 4). Figure 4 shows the likelihood values for each 3D rotation *ϕ* and candidate structures *m*. Figure 4B shows that, even when the radius of the tip apex is identical (2.5 nm), the likelihood of the structures flipped about 180 degrees have already higher likelihoods (compared to other angles), but the tip distinguishes the difference in surface geometry of each monomer and the correct angle is chosen as the maximum likelihood. On the other hand, when the radii are different from each other (Fig. 4A), the likelihood at the ground-truth angle decreases whereas the likelihoods of the 180-degrees flipped structures become competitive, resulting in the flipped structures to be selected as the maximum likelihood.

**Figure 4.**
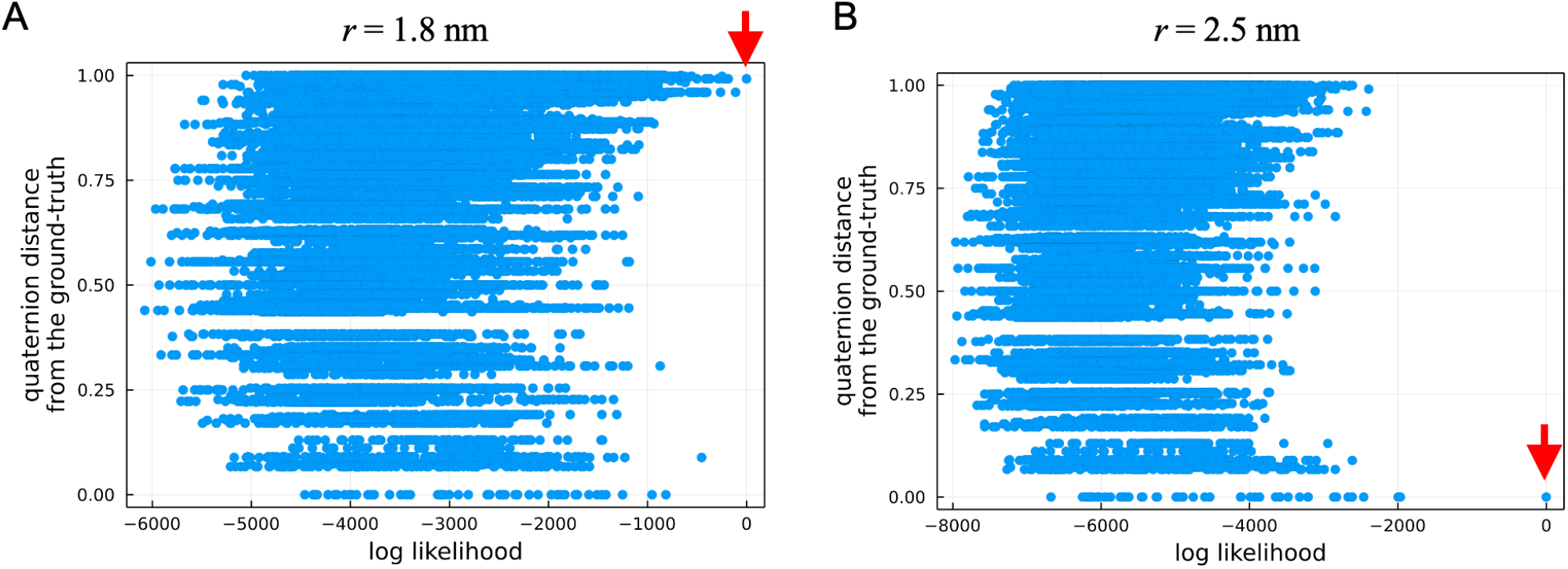
Likelihoods of all possible orientations and structures computed using the first single frame of the first set AFM data (defined by Eq. 2). The maximum likelihoods are indicated by red arrows. (A) Likelihoods computed with the tip apex radius of 1.8 nm. (B) 2.5 nm (the ground-truth).

Next, we compared the results of the frame-by-frame rigid-body fitting with those of the hidden Markov modeling (Fig. 3). We first used the ground-truth transition probabilities of MSM used for generating “experimental” AFM data. A sequence of maximum likelihood states was computed by the Viterbi algorithm. Figure 3 shows that the RMSDs of the hidden Markov modeling with the ground-truth transition matrix are overall improved compared to the RMSDs of the frame-by-frame rigid-body fitting. In particular, when the probe radius is 1.8 nm, there is a noticeable reduction in the peak around RMSD of 5.0 nm corresponding to 180-degrees flipped structures. This means that hidden Markov modeling enables a robust correct estimation of the orientation of the molecule against the choice of tip radius. As described in Eq. 3, the hidden Markov uses all frames to find the maximum likelihood sequence, not just one frame. The use of a transition matrix with rotational constraints gives penalties for large rotational flips. As a result, a fixed orientation throughout all frames is favored as the maximum likelihood sequence by a majority rule; minor orientations are not selected. This mechanism can be considered as the same principle as in the so-called ensemble learning [50]; the variance of estimation becomes large when a single discriminator is used, while the variance becomes small when multiple discriminators are combined. Note, however, when the bias getting larger due to large inconsistencies between tip radii, the majority rule cannot decrease the error as shown in Figs. 3A (1.5 nm) and 3E (3.5 nm).

To investigate how this mechanism works in multiple frames, we again examined the relation between orientations and likelihood values, changing the numbers of frames. By imposing a constraint being at a specific 3D angle *ϕ* in the Viterbi algorithm, we computed the maximum likelihood at each orientation using 1 frame, 10 frames, and 100 frames of the first set of the AFM data. Figure 5B shows the maximum likelihoods at each orientation when both radii are identical (2.5 nm). In this case, the correct orientation has already been selected as the maximum likelihood of the frame-by-frame analysis. As the number of frames increases, the difference between the largest likelihood of the correct orientation and the second largest and subsequent ones increases, indicating that the correct orientation is more likely chosen by using a greater number of frames. Figure 5A shows the results when the probe radius is not identical with the ground truth (2.5 nm and 1.8 nm). When only single frame is used, the maximum likelihood orientation is not correct, a 180 degrees flipped structure is chosen. Interestingly, as the number of frames increases, the relationship switches and the likelihood of the correct orientation begins to be chosen because the other frames have higher likelihoods for the correct orientation. As the number of frames increases, the likelihood of the correct orientation becomes more prominent. In summary, when the likelihood is calculated for only a single frame, it is probable that a wrong orientation is chosen due to the estimation error (variance), but when the number of frames is increased, the correct orientation is robustly chosen by decreasing the variance.

**Figure 5.**
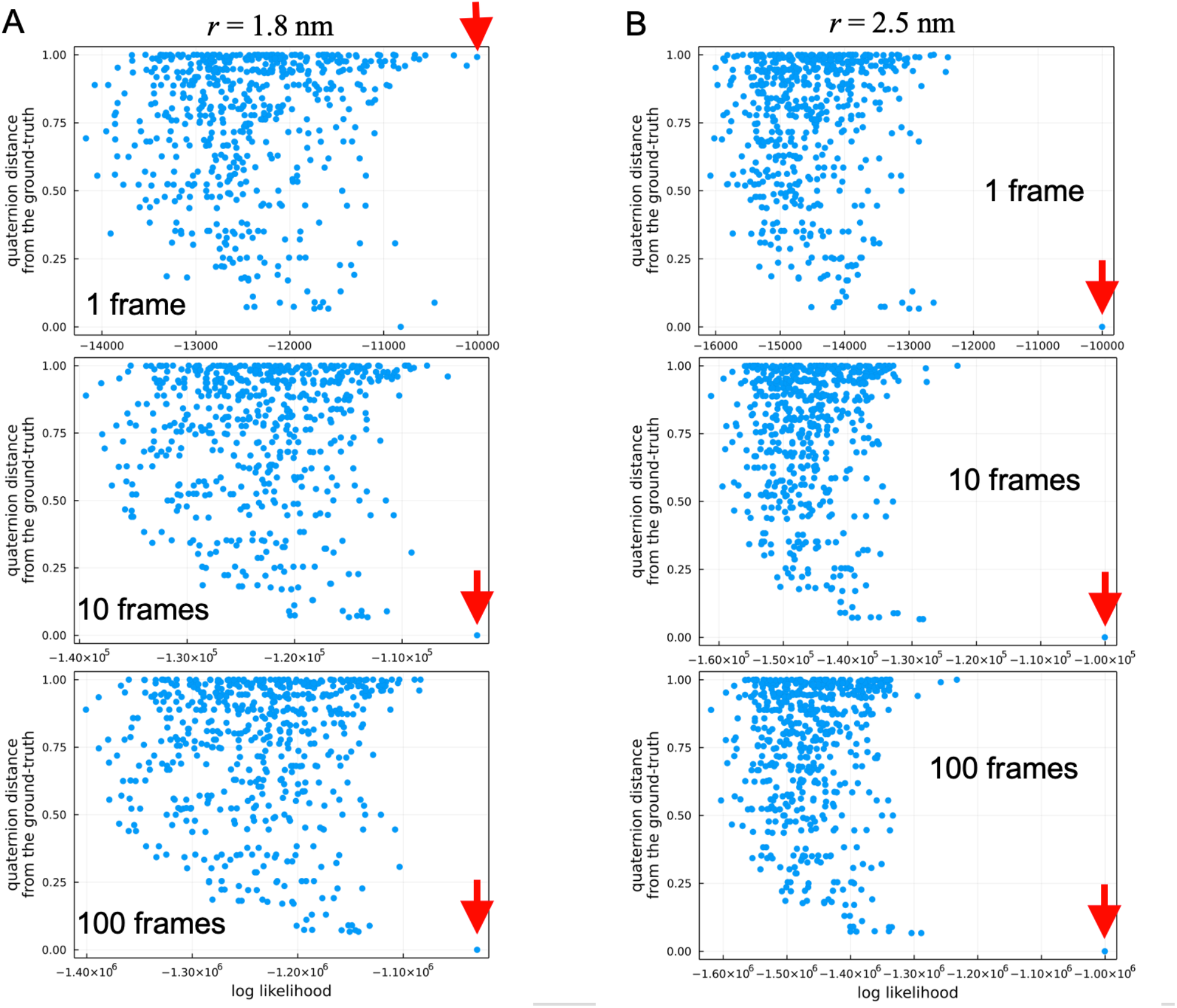
Maximum likelihood in each orientation computed using the first single frame, the first 10 frames, the first 100 frames of the first set AFM data (defined by Eq. 3). Note that only the maximum likelihood for each orientation is shown due to computational costs (calculated by the Viterbi algorithm). The maximum likelihoods among all orientations are indicated by red arrows. (A) Likelihoods computed with the tip apex radius of 1.8 nm. (B) 2.5 nm (the ground-truth).

To verify further whether the transition probabilities work to improve estimation accuracy, we computed the maximum likelihood sequence using a random matrix generated by uniform random numbers (normalized over rows) as a transition probability matrix. Interestingly, the estimation accuracy of hidden Markov modeling using the random matrix is almost identical to the results of the frame-by-frame rigid-body fitting. This is due to the fact that the mechanism of the ensemble learning does not work because the transition probabilities can no longer give penalty for large rotational flips. As a result, the likelihood of each frame is independently selected in the maximum likelihood sequence.

Finally, we performed estimations where the transition probabilities were estimated from the data without providing ground-truth transition probabilities in advance. Staring from a random matrix as the initial condition and imposing a rotation constraint, we estimated transition probabilities from the data using the Baum-Welch algorithm, and subsequently obtained the maximum likelihood sequence to verify the accuracy of the estimation. As the constraint on rotation, the probability of transitions between states whose quaternion distance is 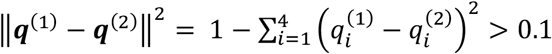 was imposed to be zero. Figure 3 shows that the estimation accuracy is improved compared with those of the rigid-body fitting, even when the transition probabilities are estimated from the data, although they are not as good as those with the ground-truth transition probabilities. This improvement in accuracy is mainly contributed by rotational constraints on the transition probabilities.

### Reconstruction of Markov state model

As calculated in the previous subsection, Hidden Markov modeling can estimate transition probabilities from the data using the Baum-Welch algorithm. Estimating transition probabilities and constructing more accurate MSM from experimental data is important topic for application studies. Here, we examined how accurately our proposed method can estimate the transition probabilities between structures using the same twin experiment data. Again, using a random matrix as the initial condition and imposing a rotation constraint, we estimated transition probabilities from the data using the Baum-Welch algorithm, Then, we marginalized the estimated transition probabilities between states (structures and rotations) to the transition probabilities between structures. After the marginalization, the probabilities were post-processed to suffice the detailed balance. Figure 6 visualizes the transition probabilities between states estimated with various tip radii. The equilibrium probability of each structure (indicated by the area of each node in Fig. 6) computed from the detailed balance.

**Figure 6.**
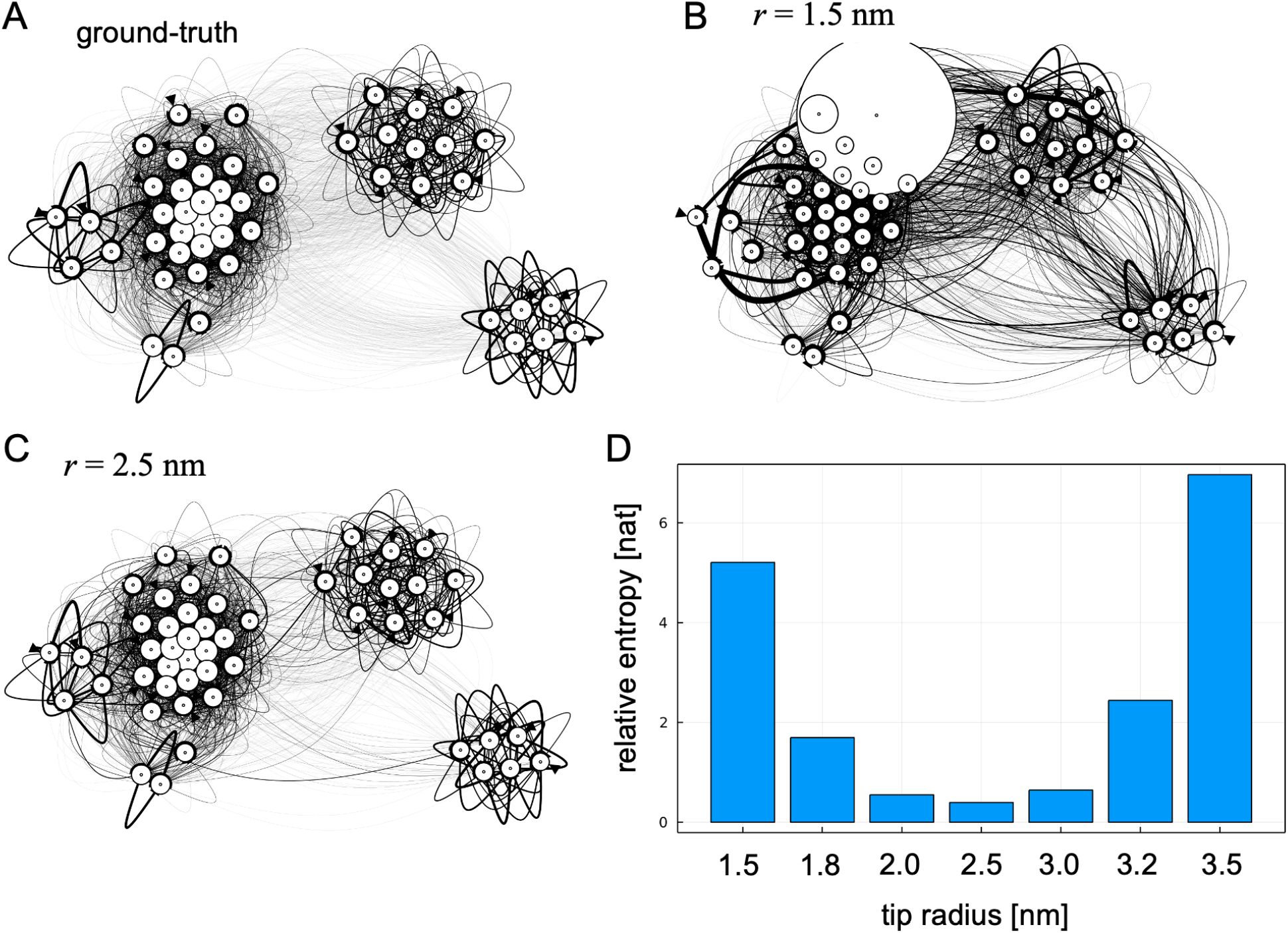
Reconstruction of MSM from atomic force microscopy data. (A) Graph representation of the ground-truth MSM used for twin experiments. (B) Reconstructed Markov state model with the tip apex radius of 1.5 nm. (C) Reconstructed MSM with the tip apex radius of 2.5 nm (the ground-truth radius). (D) Accuracies of reconstructed MSMs with various tip apex radii.

To access the accuracy of the reconstructed Markov state model, we used the relative entropy between two models [51],

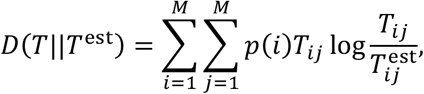

where *T*_*i j*_ is the transition probability of the ground-truth model, and *p*(*i*) is its equilibrium probability of state *i*. 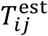 is the transition probability of the reconstructed model. The relative entropy is zero if the two models are identical and takes increasingly large values the more the two models differ. To avoid singular behavior due to zero divide, we here added a very small value (10^−10^) uniformly to 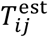.

Figure 6 shows that the estimated transition probabilities are almost the same, especially when the probe radii used for estimation are ground truth and identical (2.5 nm). In fact, as in the ground-truth MSM, CCR state is kinetic isolated from other states and the transition to this state is rare. This is true even when the probe radius is 2.0 nm, indicating that our proposed method is robust to the discrepancies in the probe geometry. According to the frame-by-frame rigid-body fitting of real AFM data (actin filament) by Nina et al. [19], the error size in the tip radii estimation from real data looks less than ± 1.0 nm. Thus, the robustness of our method would be applicable to real experimental data in the future study.

On the other hand, when 1.8 nm or 1.5 nm is used as the tip radius, the distinction between structural states becomes blurred, indicating that the kinetic isolation of CCA is significantly lost at the radius of 1.5 nm. This observation is quantified by the relative entropies between two models which takes the minimum value when two radii are identical.

### Concluding remarks

In this study, we have developed a hidden Markov modeling-based method to analyze multiple AFM images as time series data as a natural extension of the conventional frame-by-frame rigid-body fitting. In modeling AFM data, molecular states can be defined mainly by a combination of structure and rotation. Here, we have proposed a modeling that constrains transitions in rotational space to take advantage of the tendency of molecules to interact with the substrate and remain time-correlated in the rotational direction. In contrast to the conventional frame-by-frame rigid-body fitting, the rotationally constrained Hidden Markov modeling can make estimations by using the statistics of multi-frame images. By twin experiments with simulated AFM data, we have shown that the proposed hidden Markov modeling is able to estimate the molecular orientations of a taste receptor with higher accuracy.

Using the proposed method, the estimation accuracy is expected to be improved the number of frames contained in a single AFM video increases, as shown in Fig. 5. Thus, actual experimental AFM data could be modeled with higher accuracy than existing methods. However, there are several issues that must be overcome before the method applied to actual experimental data. The first issue is that a threshold for rotational constraint must be determined in advance. Although we used an arbitrary threshold for the quaternion distance in this study, it is difficult to know in advance how much the rotation of a sample molecule changes per frame. A possible prescription for this is to perform cross-correlation analysis (used in CryoEM data analysis [52]) between images of adjacent frames in AFM data, and determine a threshold from the threshold from the distribution of angular differences between adjacent frames.

Another issue is that the candidate structures for rigid-body fitting may not contain the “correct” structures corresponding to the experimental data. In this case, it is necessary to reconsider the candidate structures. A direct solution is to perform flexible fitting to AFM images using MD simulations to search for structures that are closer to the experimental images. However, flexible fitting is computationally expensive, and it is not realistic to apply it to all frames of an AFM movie. A practical way approach may be to pick up several frames at random from the HS-AFM movie, then perform flexible fitting on them to modify the candidate structures, repeating this process until the likelihood of the entire HS-AFM movie converges.

In this study, we focused mainly on orientation estimations, but in order to improve the accuracy of not only orientations but also structure estimations, it is important to use the tip shape closer to the correct one. The tip can be directly imaged by using scanning or tunnelling electron microscopy (SEM and TEM). However, since SEM and TEM provide only 2D projections of a sample, it is difficult to routinely reconstruct the 3D morphology of the tip using these approaches. Moreover, during AFM experiments, there is a possibility that the tip is partially damaged and its shape is changed. Thus, it is necessary to estimate the tip shape at the time of measurement using only AFM data. The blind tip reconstruction is a famous algorithm to estimate tip shape only from AFM image(s) [53]. The drawback of the algorithm, however, is susceptible to noise. Since HS-AFM is more prone to noise than conventional AFM, it would be necessary to develop a new method that is more robust against noise.

Although this paper described the proposed method as a natural extension of the rigid-body fitting, the method could be also used in the context of integrative structural modeling [54]. By integrating both experimental and simulated data, errors caused by model parameters in the simulations can be corrected, or experimental data can be interpreted in unprecedented details by using simulation structures. In the analysis of CryoEM density map, flexible fitting simulations has been successfully used for the refinement and interpretation of experimental data [55][56]. In the context of single-molecule measurements, Matsunaga and Sugita recently refined MSM originally constructed from simulation data using single molecule FRET measurement data to give a detailed interpretation of the experimental data [57][58]. In the same way, it would be possible to construct an accurate MSM by integrating HS-AFM and simulation data. An issue, in constructing a MSM from AFM data, is that possible dependence of structural dynamics on orientations. For some molecules, the interaction with the stage in a certain orientation may change the structural dynamics. In such cases, it is not possible to simply reduce the transition probability with respect to the orientation, as was done in this study (Fig. 6). It is necessary to check the transition probability and MSM for each orientation and to check whether the interactions with the stage cause outliers in the structural dynamics. To do this, the proposed method in this work would be useful to make reliable estimations for molecular orientations.

## Acknowledgement

We thank Shoji Takada and Sotaro Fuchigami for stimulating discussions. We also thank Toru Niina as our collision-detection code is based on his afmize. This work was supported by MEXT as “Program for Promoting Researches on the Supercomputer Fugaku” (Biomolecular dynamics in a living cell, Grant number: JPMXP1020200101 to Y.S. and Y.M.), JST CREST (Grant numbers: JPMJCR1762 to Y.M., JPMJCR13M1 to T.A.), JSPS KAKENHI (Grant numbers: 20K21380 to Y.M., 20H03195 to A.Y., 19H05645 and 21H05249 to Y.S.), and the Cooperative Research Program of “Network Joint Research Center for Materials and Devices” (to Y.M.). We used the computational resources provided by the HPCI system research project (Project ID: hp200135, hp210177, and hp220170) and those in RIKEN Hokusai “BigWaterFall”.

---

**Figure S1.**
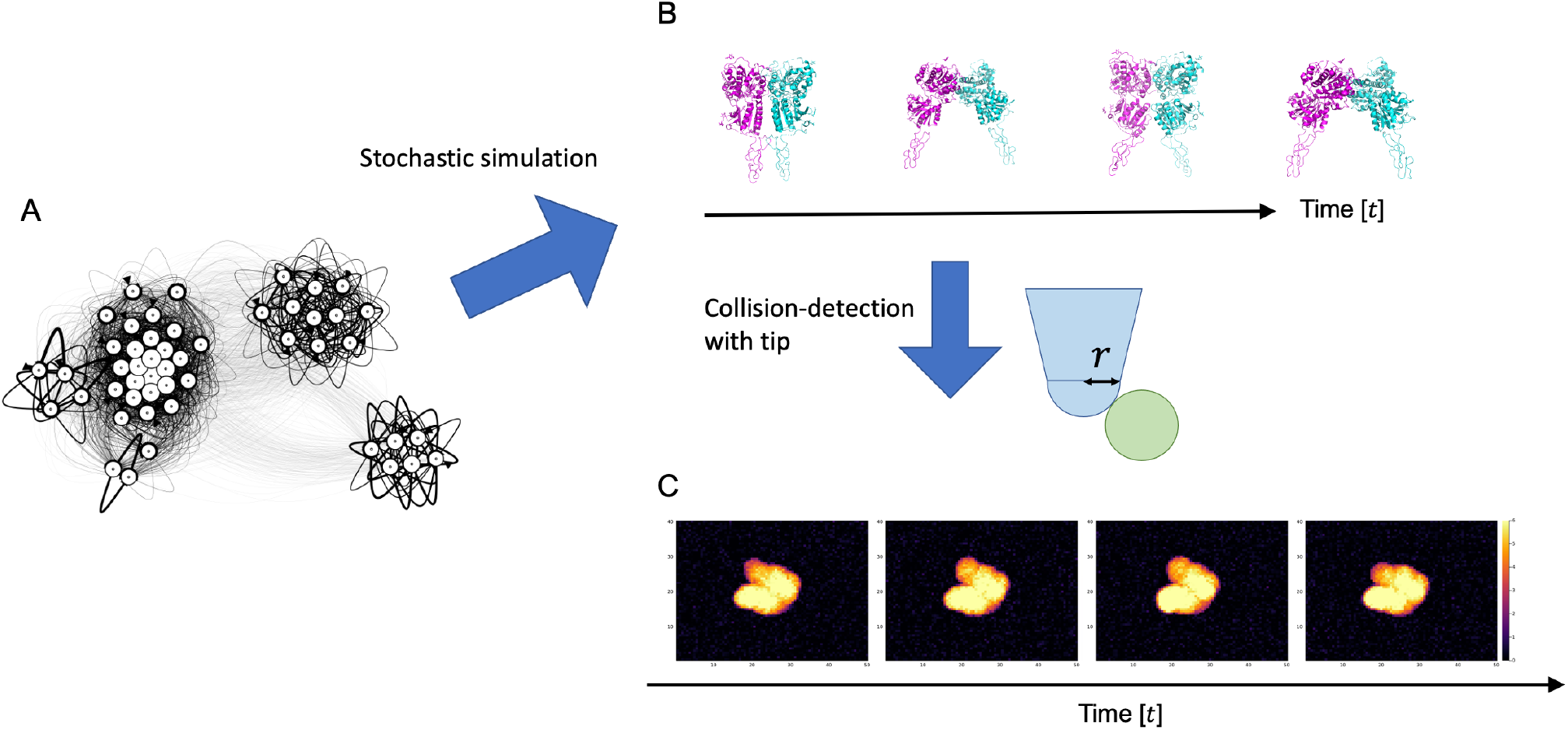
Computational protocol for generating data for twin experiment. (A) Markov state model constructed from coarse-grained molecular dynamics simulation of the taste receptor type 1. (B) A series of structures were generated by stochastic simulation of the MSM and by drawing the centroid structures of the MSM states. (C) After rotating the structures to a specific orientation, the series of AFM image were generated by the collision-detection with the tip. Finally, spatially independent Gaussian noise were added to pixels.

## Notes

### Competing Interest Statement

The authors have declared no competing interest.

## References

1. Ha T. Single-Molecule Fluorescence Resonance Energy Transfer. Methods. 2001;25: 78–86. doi:10.1006/meth.2001.1217

2. Schuler B, Eaton WA. Protein folding studied by single-molecule FRET. Current Opinion in Structural Biology. 2008;18: 16–26. doi:10.1016/j.sbi.2007.12.003

3. Moffitt JR, Chemla YR, Smith SB, Bustamante C. Recent Advances in Optical Tweezers. Annu Rev Biochem. 2008;77: 205–228. doi:10.1146/annurev.biochem.77.043007.090225

4. Binnig G, Quate CF, Gerber Ch. Atomic Force Microscope. Phys Rev Lett. 1986;56: 930–933. doi:10.1103/PhysRevLett.56.930

5. Ando T, Uchihashi T, Fukuma T. High-speed atomic force microscopy for nano-visualization of dynamic biomolecular processes. Progress in Surface Science. 2008;83: 337–437. doi:10.1016/j.progsurf.2008.09.001

6. Ando T. High-Speed Atomic Force Microscopy in Biology. Springer; 2022.

7. Kodera N, Yamamoto D, Ishikawa R, Ando T. Video imaging of walking myosin V by high-speed atomic force microscopy. Nature. 2010;468: 72–76. doi:10.1038/nature09450

8. Uchihashi T, Iino R, Ando T, Noji H. High-Speed Atomic Force Microscopy Reveals Rotary Catalysis of Rotorless F 1 -ATPase. Science. 2011;333: 755–758. doi:10.1126/science.1205510

9. Shibata M, Nishimasu H, Kodera N, Hirano S, Ando T, Uchihashi T, et al. Real-space and real-time dynamics of CRISPR-Cas9 visualized by high-speed atomic force microscopy. Nat Commun. 2017;8: 1430. doi:10.1038/s41467-017-01466-8

10. Kodera N, Noshiro D, Dora SK, Mori T, Habchi J, Blocquel D, et al. Structural and dynamics analysis of intrinsically disordered proteins by high-speed atomic force microscopy. Nat Nanotechnol. 2021;16: 181–189. doi:10.1038/s41565-020-00798-9

11. Ando T, Kodera N, Uchihashi T, Miyagi A, Nakakita R, Yamashita H, et al. High-speed Atomic Force Microscopy for Capturing Dynamic Behavior of Protein Molecules at Work. e-J Surf Sci Nanotechnol. 2005;3: 384–392. doi:10.1380/ejssnt.2005.384

12. Scheuring S, Busselez J, Lévy D. Structure of the Dimeric PufX-containing Core Complex of Rhodobacter blasticus by in Situ Atomic Force Microscopy. Journal of Biological Chemistry. 2005;280: 1426–1431. doi:10.1074/jbc.M411334200

13. Scheuring S, Boudier T, Sturgis JN. From high-resolution AFM topographs to atomic models of supramolecular assemblies. Journal of Structural Biology. 2007;159: 268–276. doi:10.1016/j.jsb.2007.01.021

14. Asakawa H, Ikegami K, Setou M, Watanabe N, Tsukada M, Fukuma T. Submolecular-Scale Imaging of α-Helices and C-Terminal Domains of Tubulins by Frequency Modulation Atomic Force Microscopy in Liquid. Biophysical Journal. 2011;101: 1270–1276. doi:10.1016/j.bpj.2011.07.020

15. Trinh M-H, Odorico M, Pique ME, Teulon J-M, Roberts VA, Ten Eyck LF, et al. Computational Reconstruction of Multidomain Proteins Using Atomic Force Microscopy Data. Structure. 2012;20: 113–120. doi:10.1016/j.str.2011.10.023

16. Chaves RC, Teulon J-M, Odorico M, Parot P, Chen SW, Pellequer J-L. Conformational dynamics of individual antibodies using computational docking and AFM: CONFORMATIONAL DYNAMICS OF IGG USING DOCKING AND AFM. J Mol Recognit. 2013;26: 596–604. doi:10.1002/jmr.2310

17. Dasgupta B, Miyashita O, Tama F. Reconstruction of low-resolution molecular structures from simulated atomic force microscopy images. Biochimica et Biophysica Acta (BBA) - General Subjects. 2020;1864: 129420. doi:10.1016/j.bbagen.2019.129420

18. Amyot R, Flechsig H. BioAFMviewer: An interactive interface for simulated AFM scanning of biomolecular structures and dynamics. Schneidman-Duhovny D, editor. PLoS Comput Biol. 2020;16: e1008444. doi:10.1371/journal.pcbi.1008444

19. Niina T, Matsunaga Y, Takada S. Rigid-body fitting to atomic force microscopy images for inferring probe shape and biomolecular structure. PLOS Computational Biology. 2021;17: e1009215. doi:10.1371/journal.pcbi.1009215

20. Amyot R, Marchesi A, Franz CM, Casuso I, Flechsig H. Simulation atomic force microscopy for atomic reconstruction of biomolecular structures from resolution-limited experimental images. PLoS Comput Biol. 2022;18: e1009970. doi:10.1371/journal.pcbi.1009970

21. Niina T, Fuchigami S, Takada S. Flexible Fitting of Biomolecular Structures to Atomic Force Microscopy Images via Biased Molecular Simulations. J Chem Theory Comput. 2020;16: 1349–1358. doi:10.1021/acs.jctc.9b00991

22. Fuchigami S, Niina T, Takada S. Particle Filter Method to Integrate High-Speed Atomic Force Microscopy Measurements with Biomolecular Simulations. J Chem Theory Comput. 2020;16: 6609–6619. doi:10.1021/acs.jctc.0c00234

23. Rabiner L, Juang B. An introduction to hidden Markov models. IEEE ASSP Magazine. 1986;3: 4–16. doi:10.1109/MASSP.1986.1165342

24. Cossio P, Hummer G. Bayesian analysis of individual electron microscopy images: Towards structures of dynamic and heterogeneous biomolecular assemblies. Journal of Structural Biology. 2013;184: 427–437. doi:10.1016/j.jsb.2013.10.006

25. Yershova A, LaValle SM, Mitchell JC. Generating Uniform Incremental Grids on SO(3) Using the Hopf Fibration. : 18.

26. Cossio P, Rohr D, Baruffa F, Rampp M, Lindenstruth V, Hummer G. BioEM: GPU-accelerated computing of Bayesian inference of electron microscopy images. Computer Physics Communications. 2017;210: 163–171. doi:10.1016/j.cpc.2016.09.014

27. Roweis S. Constrained Hidden Markov Models. : 7.

28. Nuemket N, Yasui N, Kusakabe Y, Nomura Y, Atsumi N, Akiyama S, et al. Structural basis for perception of diverse chemical substances by T1r taste receptors. Nat Commun. 2017;8: 15530. doi:10.1038/ncomms15530

29. Xu H, Staszewski L, Tang H, Adler E, Zoller M, Li X. Different functional roles of T1R subunits in the heteromeric taste receptors. Proc Natl Acad Sci USA. 2004;101: 14258–14263. doi:10.1073/pnas.0404384101

30. Husic BE, Pande VS. Markov State Models: From an Art to a Science. J Am Chem Soc. 2018;140: 2386–2396. doi:10.1021/jacs.7b12191

31. Wang W, Cao S, Zhu L, Huang X. Constructing Markov State Models to elucidate the functional conformational changes of complex biomolecules. WIREs Comput Mol Sci. 2018;8. doi:10.1002/wcms.1343

32. Chodera JD, Noé F. Markov state models of biomolecular conformational dynamics. Current Opinion in Structural Biology. 2014;25: 135–144. doi:10.1016/j.sbi.2014.04.002

33. Geng Y, Mosyak L, Kurinov I, Zuo H, Sturchler E, Cheng TC, et al. Structural mechanism of ligand activation in human calcium-sensing receptor. eLife. 2016;5: e13662. doi:10.7554/eLife.13662

34. Muto T, Tsuchiya D, Morikawa K, Jingami H. Structures of the extracellular regions of the group II/III metabotropic glutamate receptors. Proc Natl Acad Sci USA. 2007;104: 3759–3764. doi:10.1073/pnas.0611577104

35. Schlitter J, Engels M, Krüger P. Targeted molecular dynamics: A new approach for searching pathways of conformational transitions. Journal of Molecular Graphics. 1994;12: 84–89. doi:10.1016/0263-7855(94)80072-3

36. Karanicolas J, Brooks CL. The origins of asymmetry in the folding transition states of protein L and protein G. Protein Science. 2009;11: 2351–2361. doi:10.1110/ps.0205402

37. Karanicolas J, Brooks CL. Improved Gō-like Models Demonstrate the Robustness of Protein Folding Mechanisms Towards Non-native Interactions. Journal of Molecular Biology. 2003;334: 309–325. doi:10.1016/j.jmb.2003.09.047

38. Jung J, Mori T, Kobayashi C, Matsunaga Y, Yoda T, Feig M, et al. GENESIS: a hybrid-parallel and multi-scale molecular dynamics simulator with enhanced sampling algorithms for biomolecular and cellular simulations: GENESIS. WIREs Comput Mol Sci. 2015;5: 310–323. doi:10.1002/wcms.1220

39. Kobayashi C, Jung J, Matsunaga Y, Mori T, Ando T, Tamura K, et al. GENESIS 1.1: A hybrid-parallel molecular dynamics simulator with enhanced sampling algorithms on multiple computational platforms. Journal of Computational Chemistry. 2017;38: 2193–2206. doi:10.1002/jcc.24874

40. Jo S, Kim T, Iyer VG, Im W. CHARMM-GUI: A web-based graphical user interface for CHARMM. Journal of Computational Chemistry. 2008;29: 1859–1865. doi:10.1002/jcc.20945

41. Webb B, Sali A. Comparative Protein Structure Modeling Using MODELLER. Current Protocols in Bioinformatics. 2016;54. doi:10.1002/cpbi.3

42. Best RB, Zhu X, Shim J, Lopes PEM, Mittal J, Feig M, et al. Optimization of the Additive CHARMM All-Atom Protein Force Field Targeting Improved Sampling of the Backbone ϕ, ? and Side-Chain χ1 and χ2 Dihedral Angles. J Chem Theory Comput. 2012;8: 3257–3273. doi:10.1021/ct300400x

43. Huang J, MacKerell AD. CHARMM36 all-atom additive protein force field: Validation based on comparison to NMR data. Journal of Computational Chemistry. 2013;34: 2135–2145. doi:10.1002/jcc.23354

44. Jorgensen WL, Chandrasekhar J, Madura JD, Impey RW, Klein ML. Comparison of simple potential functions for simulating liquid water. J Chem Phys. 1983;79: 926–935. doi:10.1063/1.445869

45. Essmann U, Perera L, Berkowitz ML, Darden T, Lee H, Pedersen LG. A smooth particle mesh Ewald method. The Journal of Chemical Physics. 1995;103: 8577–8593. doi:10.1063/1.470117

46. Ryckaert J-P, Ciccotti G, Berendsen HJC. Numerical integration of the cartesian equations of motion of a system with constraints: molecular dynamics of n-alkanes. Journal of Computational Physics. 1977;23: 327–341. doi:10.1016/0021-9991(77)90098-5

47. Miyamoto S, Kollman PA. Settle: An analytical version of the SHAKE and RATTLE algorithm for rigid water models. Journal of Computational Chemistry. 1992;13: 952–962. doi:10.1002/jcc.540130805

48. Feig M, Karanicolas J, Brooks CL. MMTSB Tool Set: enhanced sampling and multiscale modeling methods for applications in structural biology. Journal of Molecular Graphics and Modelling. 2004;22: 377–395. doi:10.1016/j.jmgm.2003.12.005

49. Kitao A, Hirata F, Gō N. The effects of solvent on the conformation and the collective motions of protein: Normal mode analysis and molecular dynamics simulations of melittin in water and in vacuum. Chemical Physics. 1991;158: 447–472. doi:10.1016/0301-0104(91)87082-7

50. Sagi O, Rokach L. Ensemble learning: A survey. WIREs Data Mining Knowl Discov. 2018;8. doi:10.1002/widm.1249

51. Bowman GR, Ensign DL, Pande VS. Enhanced Modeling via Network Theory: Adaptive Sampling of Markov State Models. J Chem Theory Comput. 2010;6: 787–794. doi:10.1021/ct900620b

52. Tan YZ, Baldwin PR, Davis JH, Williamson JR, Potter CS, Carragher B, et al. Addressing preferred specimen orientation in single-particle cryo-EM through tilting. Nat Methods. 2017;14: 793–796. doi:10.1038/nmeth.4347

53. Villarrubia JS. Algorithms for scanned probe microscope image simulation, surface reconstruction, and tip estimation. J Res Natl Inst Stand Technol. 1997;102: 425. doi:10.6028/jres.102.030

54. van den Bedem H, Fraser JS. Integrative, dynamic structural biology at atomic resolution— it’s about time. Nat Methods. 2015;12: 307–318. doi:10.1038/nmeth.3324

55. Singharoy A, Teo I, McGreevy R, Stone JE, Zhao J, Schulten K. Molecular dynamics-based refinement and validation for sub-5 Å cryo-electron microscopy maps. eLife. 2016;5: e16105. doi:10.7554/eLife.16105

56. Mori T, Terashi G, Matsuoka D, Kihara D, Sugita Y. Efficient Flexible Fitting Refinement with Automatic Error Fixing for De Novo Structure Modeling from Cryo-EM Density Maps. J Chem Inf Model. 2021;61: 3516–3528. doi:10.1021/acs.jcim.1c00230

57. Matsunaga Y, Sugita Y. Linking time-series of single-molecule experiments with molecular dynamics simulations by machine learning. eLife. 2018;7: e32668. doi:10.7554/eLife.32668

58. Matsunaga Y, Sugita Y. Use of single-molecule time-series data for refining conformational dynamics in molecular simulations. Current Opinion in Structural Biology. 2020;61: 153–159. doi:10.1016/j.sbi.2019.12.022

